# T cell development is regulated by high fidelity replication of mitochondrial DNA

**DOI:** 10.1101/2022.12.20.521061

**Authors:** Candice B. Limper, Narda Bondah, Daphne Zhu, Alanis N. Villanueva, Uchenna K. Chukwukere, Weishan Huang, Avery August

**Affiliations:** Department of Microbiology and Immunology; Cornell Institute of Host-Microbe Interactions and Defense; Cornell Center for Immunology, Cornell University, Ithaca, NY 14853; Department of Pathobiological Sciences, School of Veterinary Medicine, Louisiana State University, Baton Rouge, LA 70803; Cornell Center for Health Equity, Cornell University, Ithaca, NY 14853

**Keywords:** DNA polymerase γ, mutator mice, T cell development, thymus, mitochondria, aging, proliferation

## Abstract

One of the most proliferative periods for T cells occurs during their development in the thymus. Increased DNA replication can result in increased DNA mutations in the nuclear genome, but also in mitochondrial genomes. A high frequency of mitochondrial DNA mutations can lead to abnormal mitochondrial function and have negative implications on human health. Furthermore, aging is accompanied by an increase in such mutations through oxidative damage and replication errors. Increased mitochondrial DNA mutations cause loss of mitochondrial protein function, and decrease energy production, substrates, and metabolites. Here we have evaluated the effect of increased mitochondrial DNA mutations on T cell development in the thymus. Using mice carrying a mutant mitochondrial DNA polymerase γ (PolG) that causes increased mitochondrial DNA mutations, we show that high fidelity replication of mitochondrial DNA is pivotal for proper T cell development. Reducing the fidelity of mitochondrial DNA replication results in a premature age-dependent reduction in the total number of CD4/CD8 double negative and double positive thymocytes. Analysis of mitochondrial density in thymocyte subpopulations suggests that this may be due to reduced proliferation in specific double negative stages. Taken together, this work suggests that T cell development is regulated by the ability of mitochondria to faithfully replicate their DNA.

## Introduction

T lymphocytes are a critical part of the adaptive immune response, needed for protection from non-self, maintenance of self-tolerance, and removing abnormal cell growth in an antigen specific manner. T cells originate from stem cells in the bone marrow, which migrate to the thymus to mature, and egress into the periphery where they can perform their effector functions. Fully functioning T cells can then proliferate and differentiate according to appropriate immune responses. One of the most proliferative periods for T cells occurs during development in the thymus. Maturation of lymphoid precursors can occur in various stages marked by CD4 and CD8 expression, i.e., CD4^-^CD8^-^ double negative (DN), CD4^+^CD8^+^ double positive (DP), CD4^+^CD8^-^ single positive or CD4^-^CD8^+^ single positive (SP). In addition, the DN stage can be further compartmentalized into four major stages based on expression of CD25 and CD44: CD44^+^CD25^-^ (DN1), CD44^+^CD25^+^ (DN2), CD44^-^CD25^+^ (DN3), and CD44^-^CD25^-^ (DN4). These different stages are defined based on different phases of the T cell receptor β (TCRβ) gene selection where the most proliferative stage is just prior to the DP stage when productive TCRβ gene arrangement occurs^1^. Such responses require synchronous metabolic support through glycolysis and oxidative phosphorylation which occur in the cytoplasm and the mitochondria, respectively^2^.

The mitochondrion is made up of around 1,000 different proteins encoded by mitochondrial and nuclear genes^3^. Approximately ~99% of the proteins in this semi-autonomous organelle are transcribed from nuclear genes. However, mitochondrial function is dependent on the mitochondrial genome. The mitochondrial genome is highly regulated to meet cellular demands and makes copies irrespective of the cell replication and in differentiating cells^4,5,6^. Access to these genes is imperative for normal mitochondrial function as they encode for 37 different mitochondrial proteins, all of which play a critical role in normal mitochondrial physiology^7^. As part of the aging process mutations accumulate in these genes, and over time these modifications of the mitochondrial genome can cause proteins to lose their functional capacity. Robust mutation repair mechanisms guard the nuclear genome of the cell^8^. In contrast, there are fewer mechanisms for correcting mitochondrial gene mutations, the most important of which is carried out by the N-terminal “proofreading” exonuclease domain of the mitochondria DNA (mtDNA) polymerase γ (PolG)^9,10^. PolG corrects mismatched mitochondrial nucleotides, which if not corrected can result in increased somatic mtDNA mutations and decreased production of energy, substrates, and metabolites by the mitochondria^10,11,12^. Immune cells, including T cells, are highly dependent on these products as they are some of the most dynamic cells in the body, being among the most proliferative cells, requiring multiple rounds of DNA replication, both nuclear and mitochondrion^13,14^.

In this study, we have evaluated the effect of increased mtDNA mutations on T cell development. Using mice carrying the *PolGD257A* mutation in the mtDNA polymerase γ (PolG) that causes increased mtDNA mutations (up to 500-fold higher mutation burden than WT mice)^18^, we show that high fidelity replication of mtDNA is pivotal for proper T cell development. Reducing the fidelity of mtDNA replication results in a premature age-dependent reduction in the total number of CD4/CD8 double positive and negative thymocytes^15^. Analysis of mitochondrial density in thymocyte subpopulations suggests that this may be due to reduced proliferation in specific double negative stages. Taken together, this work suggests that T cell development is regulated by the ability of mitochondria to faithfully replicate their DNA.

## Results

### Low fidelity mtDNA polymerase impairs thymocyte development in an age and genotype dependent manner

To determine the effects of increased mtDNA mutations on T cell development, we examined the low fidelity *PolGD257A* mouse model at different ages, analyzing age groups previously shown in the literature to represent young (1-2 months), mature (6-8 months) and old (11-13 months) mice (Figure 1A)^16,17^. We weighed the thymi and counted the total number of thymocytes from the three different age groups (young, mature, old) and genotypes; wildtype (*PolG*^+/+^), heterozygous (*PolG*^*D257A*/+^), and homozygous (*PolG^D257A/D257A^*). Consistent with previously published results, we found that there is decreased weight and total number of thymocytes (Figure 1B, C) in the *PolG^D257A/D257A^* relative to WT in the mature mice^18^. However, these differences were not observed in the young or old groups. Furthermore, we found that the *PolG^+/D257A^* mice shared the same phenotype as WT mice in all age groups, suggesting that some WT polymerase may be sufficient for the development of these cells.

**Figure 1:**
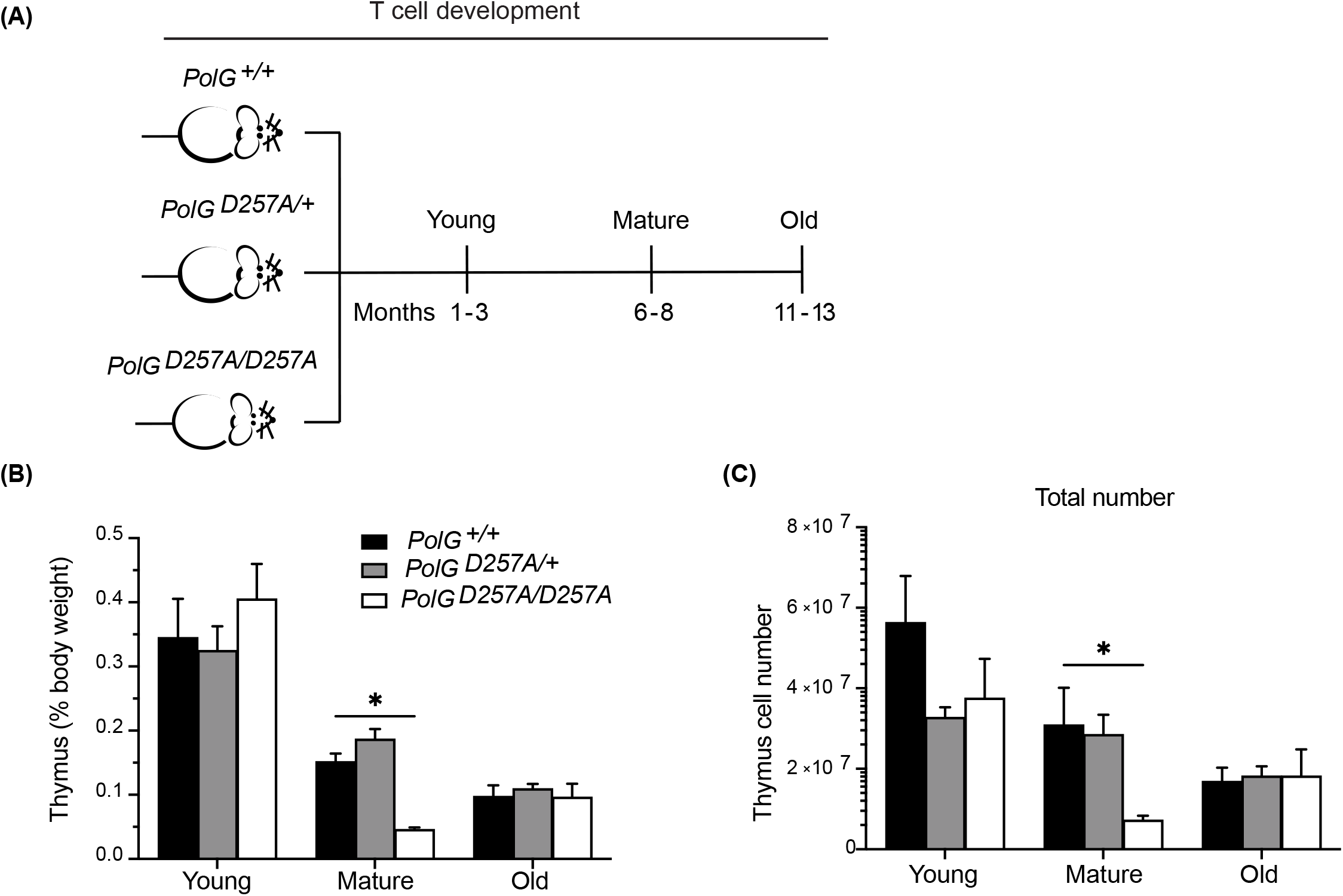
Fidelity of mtDNA replication is important for normal thymocyte numbers. **(A)** Experimental outline of *PolG^D257A/D257A^* genotypes and age groups. **(B)** Thymi from young, mature, and old *PolG^D257A/D257A^* mice were weighed and plotted as a percentage of body weight. (**C**) Thymocyte numbers from young, mature, and old *PolG^+/+^* and *PolG^+/D257A^* were determined and plotted. Data representative 11 independent experiments, n=5-24 mice per group.

We next used CD4 and CD8 to differentiate the thymocyte stages utilizing flow cytometry (Figures 2A)^19^. There was an increase in the percent of the DN population in the mature and old *PolG^D257A/D257A^* mice relative to WT mice (Figure 2B, C). Furthermore, the total cell numbers also indicated that there was fewer total number of DP and SP CD4^+^ and CD8^+^ T cells in the mature *PolG^D257A/D257A^* group relative to the WT control (Figure 2C). Taken together, this suggests that dysfunctional PolG causes a block at the DN developmental stage.

**Figure 2:**
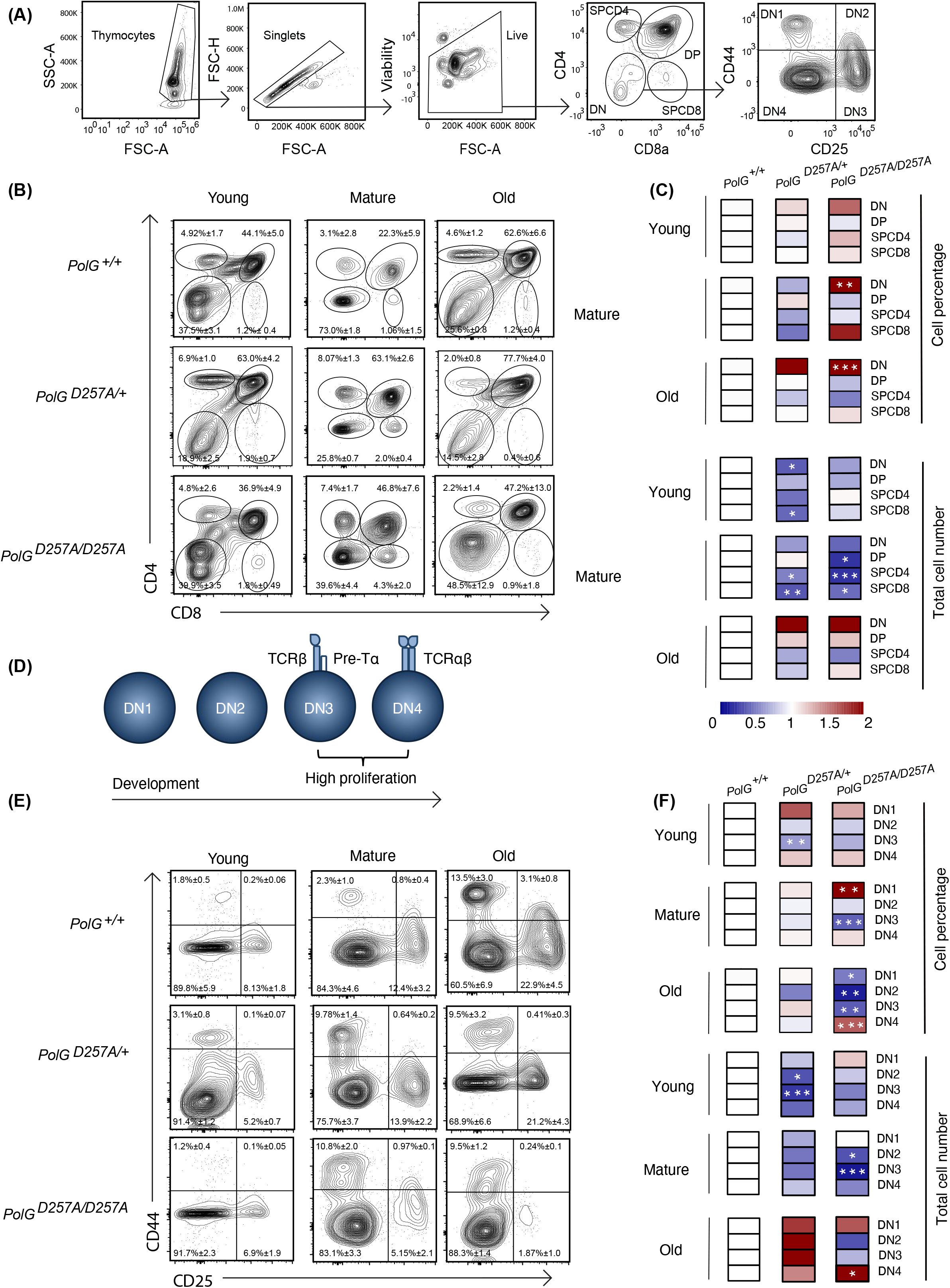
Error prone mtDNA replication impairs thymocyte development in mature *PolG^D257A/D257A^* mice. **(A)** Gating scheme used to determine the percentage (and number) of developing T cell populations. DN= double negative, DP= double positive, and SP=single positive. **(B, C)** Thymi from young, mature, and old *PolG^+/+^, PolG^+/D257A^*, and *PolG^D257A/D257A^* mice were stained for CD4 and CD8 surface markers. The gating shows DP, CD4^+^ and CD8^+^ SP thymocytes. Heatmaps are a representative quantification that are normalized by the mean of the control, where red indicates an increase and blue indicates a decrease. **(D)** Cartoon schematic showing the progression of T cell development in the DN stage. **(E, F)** Thymocytes from *PolG^+/+^, PolG^+/D257A^*, and *PolG^D257A/D257A^* mice were stained for CD25 and CD44 to differentiate DN subpopulations, and percentages of the DN1-4 stages determined. Data are displayed as mean ± standard error of mean and representative 11 independent experiments, n=7-24 mice per group, where *p< 0.05-.01, **p< 0.01-0.001, ***p<0.001 based on multiple t tests.

We sought to further investigate the DN populations with the goal of determining the stage at which these cells are being prevented from passing into the more mature stages, including the DP stage. Therefore, we investigated the four identifiable double negative populations: DN1-4, utilizing the expression patterns of CD25 and CD44 as markers (Figures 2A, D). As expected, we observed the greatest differences in the DN3 or DN4 populations, the most proliferative stages during T cell development (Figure 2E, F). We saw an increase in cell percentages of DN1, and decrease in DN3 stages, in the mature *PolG^D257A/D257A^* (Figure 2F). Furthermore, we saw a decrease in the total number of DN2 and DN3 populations in mature *PolG^D257A/D257A^* mice relative to WT mice, as well as curiously, in young *PolG^+/D257A^* mice but not in young *PolG^D257A/D257A^* mice. We also noted an increase in the number of the DN4 population in old *PolG^D257A/D257A^* mice relative to WT mice (Figure 2F). There was also what seemed like a corresponding increase in the percentage of the DN1 population, and a reduction in the percentage of DN3 population in mature *PolG^D257A/D257A^* mice relative to WT mice. The old *PolG^D257A/D257A^* mice had reduced percentages of DN1, DN2, and DN3 populations and an increase in DN4 populations. This data suggests that the presence of a low fidelity mtDNA polymerase resulted in an overall decrease in the number of DN populations and stalled T cell development, likely at the DN3 population in mature mice.

### Expression of TCR transgene partially rescues the DN population in *PolG^D257A/D257A^* mice

To interrogate the effects of the fidelity of mtDNA replication in a model of T cell development expressing a fixed TCR, we crossed *PolGD257A* mice to OT-II transgenic mice which carry rearranged TCR alpha and beta chains that are restricted to major histocompatibility complex II (MHCII) A^b^ (and recognizes the peptide 323-339 from the protein Ovalbumin)^20,21^. The presence of the already rearranged TCR accelerates the developmental stages through the DN3 when gene segments encoding the TCR alpha and beta chains would usually rearrange^22,23^. Using this transgenic T cell receptor (TgTCR) model, we tracked the T cell development in the thymus, focusing on young and mature groups given their differences in T cell development.

We collected thymi from young or mature groups and analyzed thymic weights and thymocyte number. We observed a strong concordance between the TgTCR model and the non-TgTCR *PolG^D257A/D257A^*, where we saw no significant differences in the young mice, but a reduced thymic weight of mature OTII/*PolG^D257A/D257A^* mice relative to the control (Figure 3A, B). However, when we analyzed the percentage of DN and DP populations in the OTII/*PolG^D257A/D257A^* mice relative to the control, we observed an increased overall percentage of the DN population in the OTII/*PolG^D257A/D257A^*, as was found in the non-TgTCR model, along with a decrease in the percentage of the DP population (Figure 3C). Closer examination of the DN populations revealed a trend towards an overall decrease in the percentage of the cells in the DN1-3 stages (significantly decreased in DN1), unlike what we observed in the non-TgTCR model, but an increase in the percentage of the DN4 population relative to the control (Figure 3D). This data suggests that the presence of a low fidelity mtDNA polymerase resulted in stalled T cell development at the DN3 stage in mature mice, and this is altered in the transgenic model with accelerated development through these stages.

**Figure 3:**
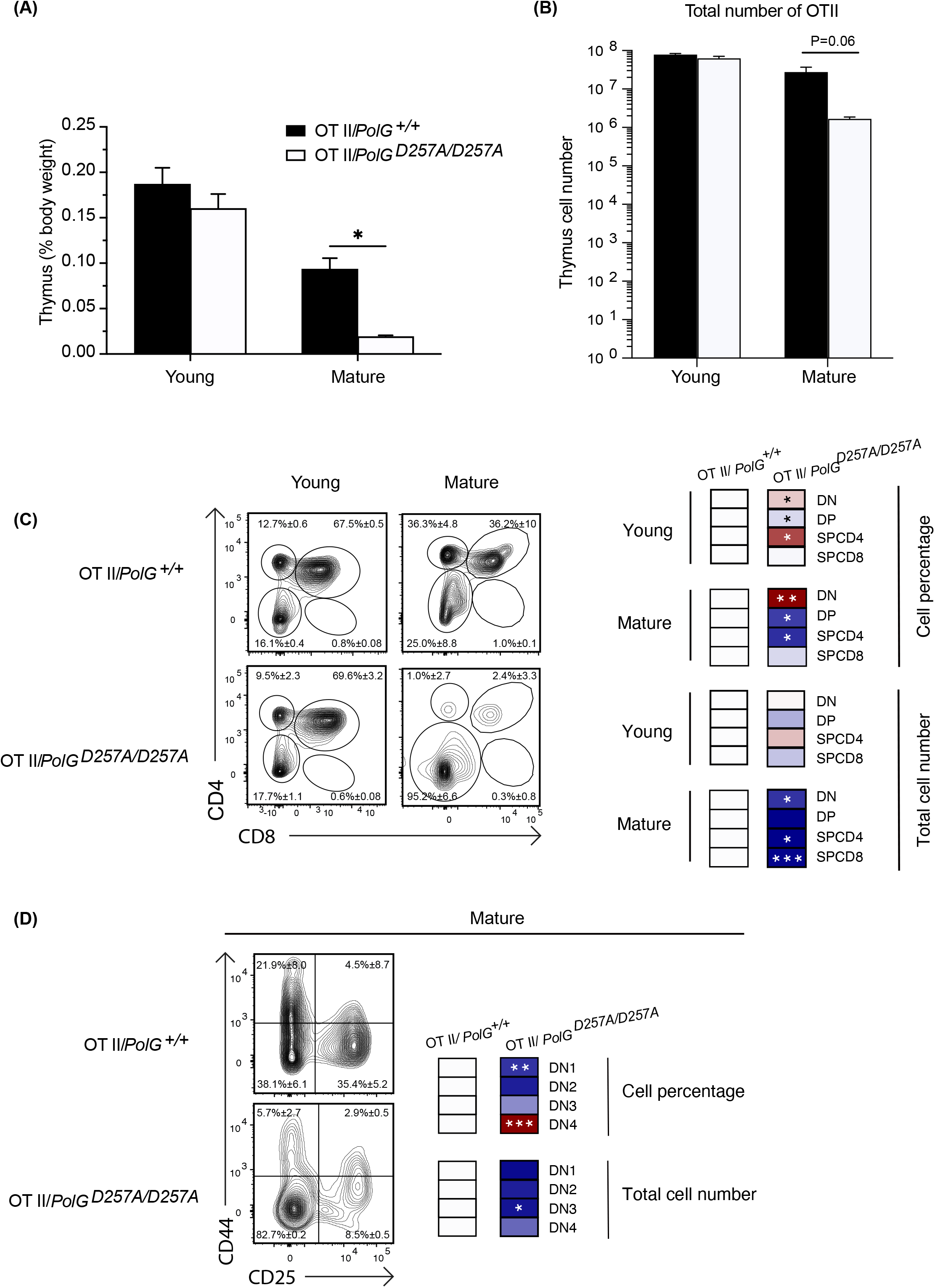
Mature OTII/*PolG^D257A7D257A^* mice have decreased thymi weight and increased DN populations relative to controls. **(A)** Thymi from mature OTII/*PolG^+/+^* OTII/*PolG^D257A/D257A^* mice were weighed and plotted as a percentage of body weight. **(B)** Thymocyte numbers from mature OTII*/PolG^+/+^ OTII/PolG^D257A/D257A^* mice were determined and plotted. p=0.06. **(C)** Thymocytes from mature OTII/*PolG^+/+^* and OTII/*PolG^D257A/D257A^* mice were stained for CD4 and CD8 and analyzed for CD4/CD8 expression. **(D)** Thymocytes from OTII/*PolG^+/+^* and OTII/*PolG^D257A/D257A^* mice were stained for CD25 and CD44 to identify DN1-4 DN subpopulations, and percentages of the DN1-4 stages determined. Heatmaps are a representative quantification that are normalized by the mean of the control, where red indicates an increase and blue indicates a decrease. Data are displayed as mean ± standard error of mean and representative of 2 independent experiments, n=3-5 mice per group, where *p< 0.05-.01, **p< 0.01-0.001, ***p<0.001 based on multiple t tests.

### Mature transgenic TCR T cells with low fidelity mtDNA polymerase have decreased mitochondrial density in DN subpopulations and exhibit less proliferation

We next wanted to explore whether the block in T cell development observed in mature OTII/ *PolG^D257A/D257A^* mice was associated with alterations in mitochondrial density within the double negative population. To investigate, we compared thymocytes from mature mice, and used a mitotracker stain, Mitotracker green, to determine mitochondrial density, gating on double negative subpopulations (DN1-4)^24^. We found that all four DN stages exhibited reduced mitochondrial density in mature OTII/*PolG^D257A/D257A^* mice (Figure 4A), with the difference increasing in the more mature stages towards DN4 (Figure 4B). Together this suggests that as thymocytes develop along the DN1-4 stages, reduced fidelity of mtDNA replication in the *PolG^D257A/D257A^* background may lead to progressively reduced mitochondrial density.

**Figure 4:**
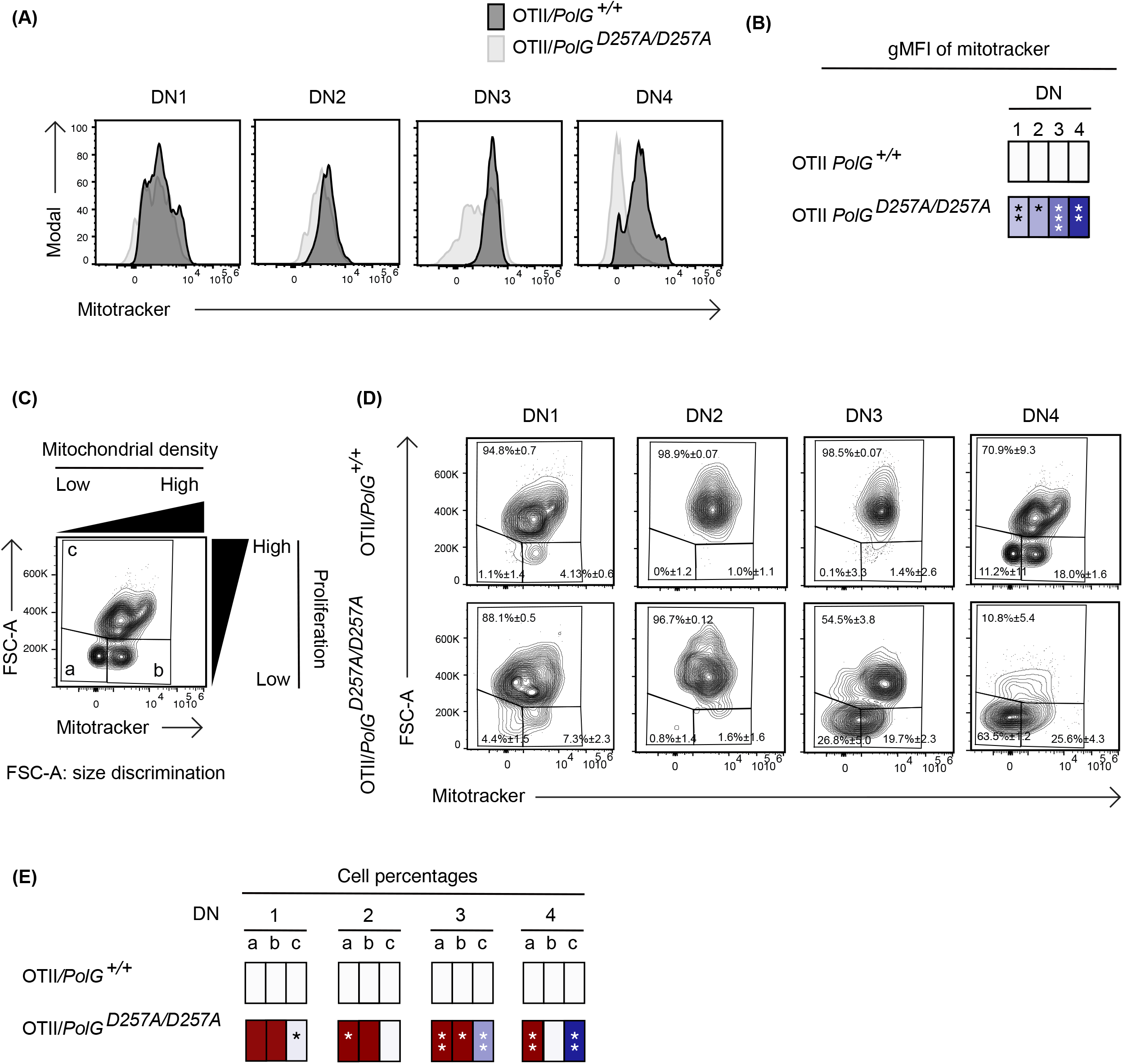
Reduced mitochondrial density and proliferation in DN thymocytes in mature OTII/*PolG*^*D257A*/D257A^ mice. **(A)** Thymocytes from OTII/*PolG^+/+^* and OTII*/PolG^D257A/D257A^* mice were stained for mitochondria, along with CD25 and CD44. Histograms depict mitochondria density in gated double negative subpopulations: DN1, DN2, DN3, and DN4. **(B)** Geometric mean fluorescence intensity (gMFI) of mitochondria density staining in DN1-4 thymocytes from OTII*/PolG^+/^*^+^ and OTII/*PolG*^*D257A*/D257A^ mice. **(C)** Illustration of gating strategy used to discriminate larger proliferating cells and mitochondrial density and **(D,E)** frequencies of size (FSC-A) and mitochondrial density (mitotracker stain) in DN1-4 populations as indicated. Heatmaps are a representative quantification that are normalized by the mean of the control, where red indicates an increase and blue indicates a decrease. Data are displayed as mean ± standard error of mean and representative of 1 independent experiment, n=3 mice, where *p< 0.05-.01, **p< 0.01-0.001, ***p<0.001 based on multiple t tests.

Since these DN populations are known to be highly proliferative, we wanted to explore whether decreased mitochondria negatively impact this highly proliferative stage of T cell development in the mature OTII/*PolG^D257A/D257A^* mice^25,26,27^. Figure 4D shows an analysis that determines proliferation (based on cell size using FSC-A, since the larger cell size is associated with cell cycle progression)^28,27^, where populations *a* and *b* are low proliferation populations, and population *c* is high (Figure 4C). The proportion of large (proliferating) DN3 and DN4 cells in OTII/*PolG^D257A/D257A^* thymocytes (as determined by the increase in cell size) was drastically decreased relative to the control (Figure 4D). Additionally, there was a decrease in the mitochondrial density (shown as an increase in the cells falling in gate *a*) in the OTII/*PolG^D257A/D257A^* DN3, and again more drastically in DN4 populations relative to OTII/*PolG*^+/+^ (Figure 4D, E).

### Mature OTII/*PolG^D257A/D257A^* T cells have decreased mitochondrial density and proliferation at the SP CD4^+^ T cell stage

Lastly, we wanted to determine whether the aberrant effects observed in the mature OTII/*PolG^D257A/D257A^* continued through the rest of T cell development in the thymus. To determine this, we looked at more mature DP and SPCD4 T cell populations in the thymus. We saw no difference in the mitochondrial density (gates *a* and *b*), but there was a decrease in the proliferating cell (larger cells in gate *c*) in the OTII/*PolG^D257A/D257A^* relative to the control (Figure 5A). There was also a decrease in the mitochondrial density in the OTII/*PolG^D257A/D257A^* SPCD4^+^ T cells relative to the control, although no observed difference in proliferating (larger) cells between OTII/*PolG*^+/+^ and OTII/*PolG^D257A/D257A^* (Figure 5B).

**Figure 5:**
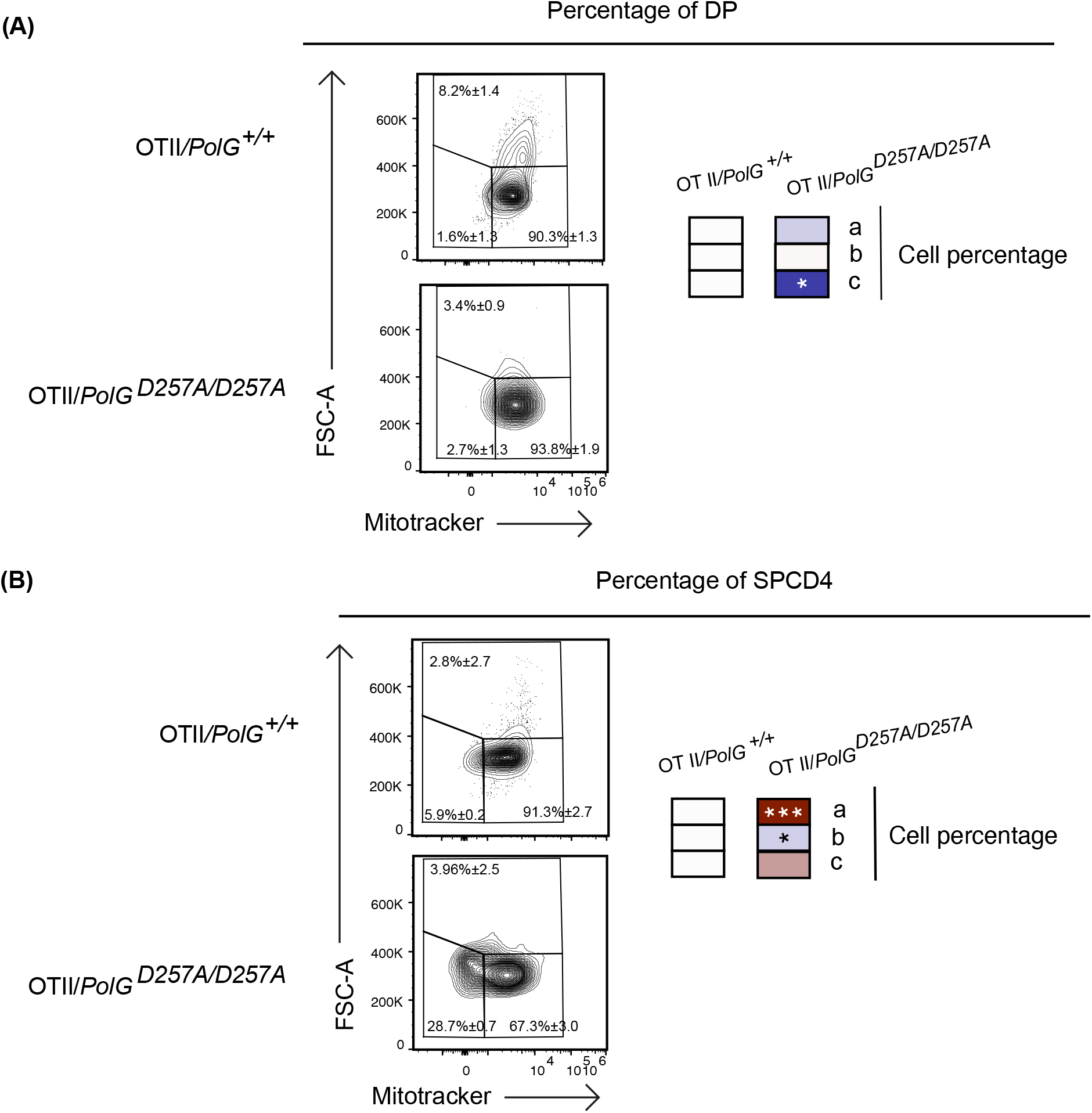
Single positive CD4^+^ T cells in mature OTII/*PolG*^*D257A*/D257A^ mice have decreased mitochondrial density. **(A)** Thymocytes from OTII/*PolG*^+/+^ and OTII/*PolG^D257A/D257A^* mice were stained for mitotracker, along with CD4 and CD8, and the frequencies of size (FSC-A). The histograms represent mitochondrial density (mitotracker stain) in CD4/CD8 DP **(B)** or CD4 SP population plotted. Data are displayed as mean ± standard error of mean and representative of 1 independent experiment, n=3 mice, where *p< 0.05-.01, **p< 0.01-0.001, ***p<0.001 based on multiple t tests.

## Discussion

The role of mitochondria in T cell development has only recently been intensely investigated, and understanding the effects of mtDNA mutations on this process is even more limited^2^. Here, we have explored the effects of reduced mtDNA replication fidelity, which leads to increase in mtDNA mutations in mice carrying the mutant *PolGD257A*, on T cell development. We found that reducing the fidelity of mtDNA replication results in premature age-dependent reduction in the total number of CD4/CD8 double positive and negative thymocytes. This is likely due to reduced proliferation in the highly proliferative double negative stages due to reduced mitochondrial density and the accompanying effect on mitochondrial products needed for appropriate function. Taken together, this work suggests that T cell development is regulated by the ability of mitochondria to faithfully replicate their DNA.

We examined the importance of the fidelity of the mtDNA replication in T cell development and the significance of this process in an age dependent manner, given that over time mtDNA mutations accumulate naturally in WT mice and at an accelerated rate (>500 fold) in *PolGD257A* mice^18^. Others have previously shown that homozygous mutations in the exonuclease region of *PolGD257A* result in premature aging phenotypes, including the thymus, in 9-13 month old mice^18^. Interestingly, 3 month old mice have an increase in apoptosis in total thymi, suggesting that there might be fewer thymocytes because of increased cell death^18^. Considering the age gap between 3 and 9 month old mice, we sought to determine whether these differences are detectable between these ages, and whether these findings depend on the dose of the PolGD275A mutant. Similar to previous findings, we saw decreased weight and total cell number of thymocytes in the mature *PolG^D257A/D257A^* mice^18,29^. That this observation was not detected in the heterozygous group suggests that one functional copy of the PolG gene may be sufficient to otherwise ameliorate negative effects.

We further investigated the effects on various T cell developmental stages. The DN stage includes TCR β chain-selection, survival, differentiation, proliferation, and allelic exclusion at the TCRβ locus^30^. We predicted that we would see an increase in the DN stage in the mature *PolG^D257A/D257A^* mice relative to the control because there would be higher mutations due to the high proliferation rates, and the findings supported this prediction. There are different factors that distinguish progression through the DN stage, discriminated between DN1-4 stages. DN1-2 or early thymic precursors are the earliest clearly identifiable intrathymic stages^23^. We observed an increase in the DN1 population in the mature *PolG^D257A/D257A^* mice relative to the controls. This suggests that there may be a developmental stall in the DN1 stage, thus reducing progression to the subsequent stages and affecting further T cell development.

T cell commitment occurs at the end of the DN2 prior to entry into the DN3, and during this stage they traverse a significant proliferative expansion^31^. We observed a decrease in the cell number and percentage of DN3 population in the *PolG^D257A/D257A^* mice, suggesting that the fidelity of mtDNA replication significantly affects the highly proliferative cells in the DN3 stage. A possible reason for the reduction in the number of these cells is increased cell death, although this requires further investigation. This finding highlights the importance of functional mitochondria and mitochondrial products in this highly proliferative stage, which may be more sensitive to higher mtDNA mutational burden in the *PolG^D257A/D257A^* mice.

The DP stage, which follows the highly proliferative DN stage, is where the CD4 and CD8 co-receptors are upregulated and expression of the rearranged TCRα and β chains occur^30^. Following which, T cells go through positive selection where the TCRs are tested for having appropriate avidity for the MHC/peptide. These stages do not require high proliferation like the DN stage. Therefore, we predicted that we would not see a difference in the percentage of these cells, but a decrease in the cell number in the *PolG^D257A/D257A^* mice relative to the control, and this conclusion aligned with the data.

We also examined the effect of increased mtDNA mutations on T cell development in OTII/*PolG^D257A/D257A^* transgenic mice. The rationale for our analysis of OTII/*PolG^D257A/D257A^* mice was that the presence of the already rearranged TCR accelerates the progress through the DN developmental stages, when gene segments would normally rearrange and encode the TCR alpha and beta chains. Unlike the non-transgenic mice, T cell development in the transgenic mice do not have to go through TCR gene rearrangement in the same manner as non-transgenic mice, since unlike the non-transgenic mice, T cells in the transgenic mice do not have to go through TCR VDJ recombination at the alpha and beta loci during their development^32^. Thus, if the increased proliferation during these stages was inordinately affected by the low fidelity mtDNA replication, then these stages could potentially be less affected in these transgenic thymocytes. Notably, we did not see the same trend of increased numbers of thymocytes in the DN stages in the OTII/*PolG^D257A/D257A^* mice. However, we found that there was a block later in the DN stage, with an increase in the percentage of DN4 in the mature OTII/*PolG^D257A/D257A^* mice relative to the control.

We explored potential mechanisms for these observed defects in T cell development in the mutant mice by analysis of mitochondrial density and cell size, as a proxy for cell division. Abnormal mtDNA replication can lead to altered expression of mitochondrial proteins which are important for the electron transport chain and can cause decreased oxidative phosphorylation and activate mitochondrial degradation through mitophagy. If there are fewer mitochondria to make appropriate substrates for cell replication, this can alter metabolism and cause deleterious effects to cell proliferation^2^. Further analysis of the mitochondrial density indicated that OTII/*PolG^D257A/D257A^* DN1-4 populations are decreased compared to the controls. Furthermore, there is a notable reduced proliferation in DN3 and DN4 populations in the OTII/*PolG^D257A/D257A^* relative to WT. This trend appeared to increase over differentiation stage, where the largest differences between mitochondrial density and proliferation were observed at the DN4 stage. This could be due to a larger buildup of mtDNA mutations relative to the control at this stage. Furthermore, it is possible that increased mtDNA mutation could result in decreased mitochondrial function and increased mitophagy. One major limitation to our findings is that it is unclear whether they are influenced by non-T cell dependent factors given that *PolG^D257A/D257A^* and OTII/*PolG^D257A/D257A^* models are globally mutated. Therefore, future experiments will need to be conducted to determine whether our observations are due to T cell intrinsic effects.

There are drugs with reported side effects of decreased mtDNA content and mitochondrial functional impairment similar to what is observed in the PolG mouse model^33,34^. One common observation is mitochondrial toxicity seen with the use of reverse transcriptase inhibitors which can resemble purines (adenosine and guanosine) and pyrimidines (cytidine, thymine, and uridine), known as nucleoside-analogue reverse-transcriptase inhibitor. These inhibitors are widely utilized as an antiviral treatment with patients who have viral infections^35^. It is currently unclear whether such drugs induce similar alterations as we observe in the PolG mutant mice.

Altogether our data revealed that the fidelity of mtDNA replication is critical to T cell development, affecting proliferation in specific DN stages. This reduced proliferation and block in the DN stages may be due to decreased mitochondrial density, resulting in reduced proliferation and development of T cells. This study contributes to our understanding of PolG and the fidelity of mtDNA replication in T cell development, providing new insight into how mitochondria affect this process.

## Materials and Methods

### Mice

All mice were maintained and housed in specific pathogen-free facilities and experiments were performed in accordance with protocols that were approved by the Institutional Animal Care and Use Committee at Cornell University. *PolG^D257A/D257A^* mice have been previously described and obtained from the Jax labs^18^. Male *PolG*^+/D257A^ and *PolG^D257A/D257A^* were backcrossed with WT C57BL/6J, mice, to generate male and female heterozygous and homozygous littermates which were then crossed and used for this study. The presence of *PolGD257A* knock-in mutation was determined by PCR using the following primers in genotyping (5’ to 3’; reverse common: AGT CCT GCG CCA ACA CAG; wildtype forward: GCT TTG CTT GAT CTC TGC TC; mutant forward: ACG AAG TTA TTA GGT CCC TCG AC). OTII transgenic mice were also obtained from the Jax labs and crossed with *PolG^D257A/D257A^* and then the OTII/*PolG^D257A/+^* were crossed with OTII/*PolG^D257A/+^* and the subsequent generation were used for experiments.

### Antibodies for flow cytometry

The following antibodies were used for flow cytometry analysis: PerCP-Cy5.5-anti-CD25, PE-anti-CD24, CD44-FITC, PE-CF594-anti-CD4 (BD Biosciences, Inc.), PacificBlue-anti-CD4, PacificBlue-anti-CD8a, eF506-viability dye (eBioscience Inc.), and AF700-anti-CD8a, Pecy7-anti-TCRβ (BioLegend). Mitotracker green (Cell Signaling Technology) was used to detect mitochondrial density.

### Flow cytometry

Organs were mechanically dissociated through 70 μM screen placed in complete RPMI media while on ice. For surface staining, cells were stained, washed, and all flow cytometry analysis was conducted in the Cornell University Flow Cytometry Core Facility using the Thermo Fisher Attune NxT and data was analyzed in FlowJo (Tree Star, Ashland, OR) where all data were performed gating on doublet excluded viable cells.

### Data extraction and statistical analysis

Flow cytometry data was extracted and analyzed using code through RStudio 2022.07.1 Build 554 (GitHub repository TMPierpont/tmisBasic/Heatmap/R/timsDataLoader.R 2022). This code was used RStudio to export percentages from FlowJo analysis and to calculate total cell numbers. The flow cytometry gates show average cell percentages, and heatmaps are a representative quantification that are normalized by the mean of the control. Statistical analyses were carried out utilizing Prism 9 (GraphPad, San Diego CA) and RStudio. Statistical significance was set at p<0.05, and all data are reported as a mean, plus and minus standard error of the mean (SEM). Age and genotype were compared by multiple t-tests.

## Acknowledgements

We thank the Cornell animal care facilities for their animal care, Amie Redko for maintaining the breeder colony and Ling Zhang for genotyping. The following Cornell University core facilities have also made this research possible: the Institute of Biotechnology for collecting the data and Statistical Consulting Unit for assistance with data analysis. Lastly, we thank members of the August lab for comments and suggestions.

## Author Contributions

AA conceived the project, supervised experiments, and analyzed data. CBL, NB, DZ, UKC, ANV, and WH performed experiments. CBL analyzed the data. AA and CBL wrote the manuscript, and all authors approved the submitted version.

